# Intrinsic Cell-type Selectivity and Inter-neuronal Connectivity Alteration by Transcranial Focused Ultrasound

**DOI:** 10.1101/576066

**Authors:** Kai Yu, Xiaodan Niu, Esther Krook-Magnuson, Bin He

## Abstract

Transcranial focused ultrasound (tFUS) is a promising neuromodulation technique, but its mechanisms remain unclear. We investigate the effect of tFUS stimulation on different neuron types and synaptic connectivity in *in vivo* anesthetized rodent brains. Single units were separated into regular-spiking and fast-spiking units based on their extracellular spike shapes, further validated in transgenic optogenetic mice models of light-excitable excitatory and inhibitory neurons. For the first time, we show that excitatory neurons are significantly less responsive to low ultrasound pulse repetition frequencies (UPRFs), whereas the spike rates of inhibitory neurons do not change significantly across all UPRF levels. Our results suggest that we can preferentially target specific neuron types noninvasively by altering the tFUS UPRF. We also report *in vivo* observation of long-term synaptic connectivity changes induced by noninvasive tFUS in rats. This finding suggests tFUS can be used to encode temporally dependent stimulation paradigms into neural circuits and non-invasively elicit long-term changes in synaptic connectivity.

## INTRODUCTION

Neuromodulation is to interven with the nervous system to treat and help improve the quality of life of subjects suffering from neurological disorders. For decades, a myriad of brain neuromodulatory approaches, such as deep brain stimulation (DBS) (Ashkan et al., 2017), transcranial magnetic stimulation (TMS) (Barker et al., 1985; Kobayashi and Pascual-Leone, 2003), transcranial current stimulation (tCS) (Horvath et al., 2015; Nitsche and Paulus, 2000), transcranial focused ultrasound (tFUS) (Lee et al., 2016b; Legon et al., 2014; Tufail et al., 2010), transcranial static magnetic field stimulation (tSMS) (Oliviero et al., 2011), optogenetics (Boyden et al., 2005; Chen et al., 2018; Deisseroth and Hegemann, 2017), designer receptors exclusively activated by designer drugs (DREADDS) etc., have been developed in order to modulate and study the brain. Among these methods, optogenetics receives considerable attention for its capacity to selectively stimulate distinct cell-types (Packer et al., 2013; Rajasethupathy et al., 2016) with high spatial and temporal resolution (Boyden et al., 2005). However, optogenetics heavily relies on methods such as transgenic approaches, viral vector transfection or nanoparticle injection for deep brain application (Chen et al., 2018), which pose practical challenges for translation in human clinical utility (Delbeke et al., 2017). In contrast, non-invasive methods such as TMS and tCS are readily translated to clinical utility, but are challenged to achieve highly spatial focus and deep penetration.

As a promising new technique, low-intensity tFUS can be applied in many neuromodulation applications due to its high spatial focality (compared to TMS and tCS (Polania et al., 2018)) and its non-invasive nature (Naor et al., 2016). During tFUS neuromodulation, pulsed mechanical energy is transmitted though the skull with high spatial selectivity (Legon et al., 2014), which can be steered (Haritonova et al., 2015) and utilized to elicit activation or inhibition through parameter tuning (King et al., 2013; Ye et al., 2016). Pilot studies have investigated the neural effects of ultrasound parameters, such as ultrasound fundamental frequencies (UFF), intensities (UI), durations (UD), duty cycles (UDC), pulse repetition frequencies (UPRF), etc. Besides a few human studies (Hameroff et al., 2013; Lee et al., 2016b; Legon et al., 2014), animal models, such as worms (Ibsen et al., 2015; Kubanek et al., 2018; Zhou et al., 2017), rodents (Tufail et al., 2010; Ye et al., 2016; Yu et al., 2016), rabbits (Yoo et al., 2011), swine (Dallapiazza et al., 2018), and monkeys (Deffieux et al., 2013; Folloni et al., 2019), have been utilized to investigate the effects of ultrasonic parameters and acoustic-induced effects. tFUS have been observed to induce behavioral changes, e.g. motor responses (King et al., 2013; Mehic et al., 2014), electrophysiological responses, e.g. electromyography (EMG) (King et al., 2014; Ye et al., 2016), electroencephalography (EEG) (Darvas et al., 2016; Yu et al., 2016), local field potentials (LFPs) (Tufail et al., 2010), and multiunit activities (MUAs) (Tufail et al., 2010), with high *in vivo* temporal / spatial measurement fidelity, or neurovascular activities, e.g. blood-oxygenation-level-dependent (BOLD) signal (Folloni et al., 2019; Verhagen et al., 2019; Yang et al., 2018; Yoo et al., 2011), etc. To further achieve selectivity in stimulating brain circuits or even among cellular populations, focused ultrasound is employed in combination with specific-neuromodulatory-drug-laden nanoparticles (Airan et al., 2017), cell-specific expression of ultrasound sensitizing ion channels (Ibsen et al., 2015) or acoustically distinct reporter genes in microorganisms (Bourdeau et al., 2018). Based on previous studies, it is evident that distinct tFUS parameters can exhibit unique stimulation effects. So far, none have explored the intrinsic effects of the wide range of ultrasound parameters on specific neuron subpopulations.

Exploring ultrasound’s intrinsic cell-type selectivity may pave the way for the translation of tFUS as an effective non-invasive modulation tool for brain conditioning and elucidating neural mechanisms. A unifying theoretical framework was formulated in order to map the acoustic parameters that lead to neuronal activation/suppression effects through cortical stimulation results based on their neuronal intramembrane cavitation excitation (NICE) model (Plaksin et al., 2016; Plaksin et al., 2014). While further validations are necessary, certain cell-type-selective effects are predicted by this model when T-type calcium channels, existing in both cardiac and central nervous systems, are added in the neuronal modeling (Plaksin et al., 2016). Inspired by this work and the increasing number of studies demonstrating ion channel dynamics as the mechanism of tFUS stimulation (Bystritsky et al., 2011; Tyler, 2011; Tyler et al., 2008), we investigate the cell-type dependent effects of tFUS stimulation through extracellular recordings in *in vivo* rodent brains. As shown in Fig. 1, we hypothesize that different types of neurons will have distinct response profiles to the dynamic acoustic radiation forces exerted by the UPRF of tFUS. The differences observed between different cell types may be contributed by the types and relative distributions of ion channels in each cell type and/or the distinct shape and orientation of the axonal and dendritic arbors.

**Figure 1.**
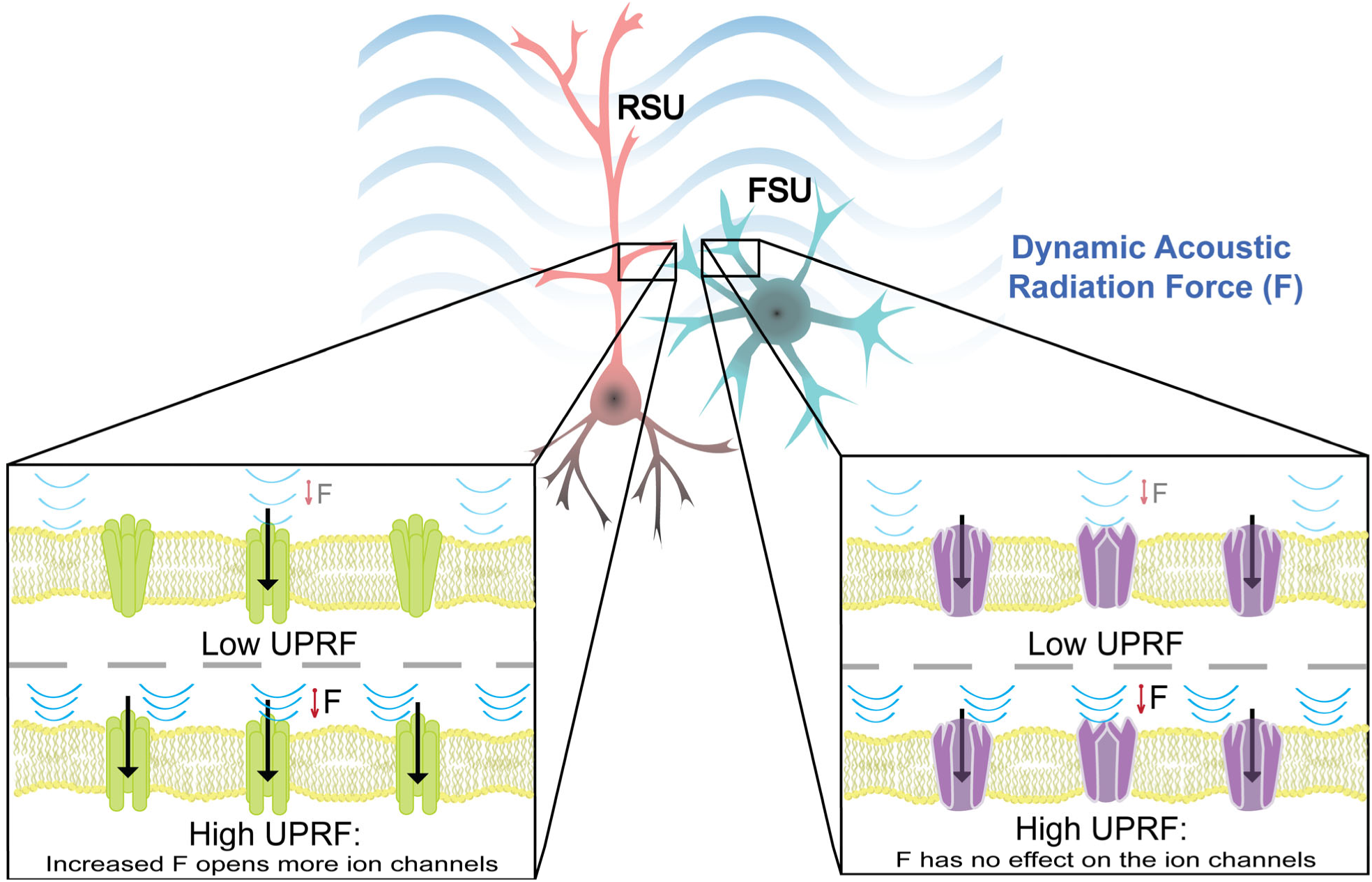
Conceptual Diagram Illustrates Hypothetical Differences of Types of Neurons in Responding to the Dynamic Acoustic Radiation Force (ARF). A hypothesized mechanism for observed difference in response to tFUS in distinct cell types. The dynamic acoustic radiation force is induced by the ultrasound pulse repetition frequency (UPRF). The regular spiking units (RSU) exhibit more sensitivity to the ARF change than the fast spiking units (FSU) do.

Beyond investigations on the short-term intrinsic effects of UPRF on neuron subtypes, we also explore the ability to use tFUS parameters for encoding frequency specific information into the brain for long-term effects. One step further than studying ultrasound effects on single cortical neuronal activities, we investigate potential sustained therapeutic effects of acoustic energy on inter-neuronal synaptic connectivity at deep brain. tFUS targeting the deep brain were demonstrated recently (Dallapiazza et al., 2018; Legon et al., 2018), and non-transient neuromodulatory effects were also observed in anesthetized rat (Niu et al., 2017; Yoo et al., 2017) and monkey (Verhagen et al., 2019) models. These are early evidences of ultrasound-induced neuroplasticity, a concept which we further explored in the well characterized system of hippocampal long-term potentiation (LTP) and depression (LTD) in rats. The innovation of this work is that instead of inducing long term effects through the passive application of prolonged and intensified stimulations used in the previous studies, we further investigate eliciting sustained effects through the active encoding of neurologically relevant frequency to the hippocampus.

LTP and LTD have been widely studied in neuroscience as a mechanism of memory storage and emotion processing in the hippocampus (Min et al., 1998). The LTP literature has shown that brief high frequency electrical stimulation trains applied at the Schaffer collaterals increase the EPSP slope amplitudes seen in the CA1 pyramidal cells, as well as in the medial perforant path to the dentate gyrus (Dudek and Bear, 1993; Dudek and Bear, 1992; Purves D, 2001). When stimulated with low frequency electrical stimulation, LTD (decrease in EPSP strength) can be observed (Barrionuevo et al., 1980; Dudek and Bear, 1992). This illustrates the immense ability of the brain to use frequency encoding for information storage. Similar to electrical stimulation, we hypothesize that tFUS has the ability to increase the postsynaptic potential and when applied at a comparable frequency as tetanus electrical stimulation, induce LTP or LTD.

Throughout our studies, we were attentive to potential biases and confounding variables in our data. Recently, two published companion studies brought to attention the possibility of auditory confounding effects of ultrasound neuromodulation (Guo et al. 2018, Sato et al. 2018). The findings of the companion studies call into attention confounding cortical activations that may arrive due to ultrasound mechanical coupling with the rodent skull. To control for these effects in our set up, we have conducted control studies in deafened rats to ascertain that local activations are present in our setup without auditory percepts (Niu et al., 2018). On the naive rats in this study, we also conducted more sham studies to control for potential false positive findings.

## RESULTS

### Characterizing tFUS Stimulation and Setup

The *in vivo* experimental setup and ultrasound temporal and spatial profiles are illustrated in Fig. 2. As presented (Fig. 2A), pulsed tFUS is first generated by a single-element transducer, guided to a scalp location over the left somatosensory cortex through a mounted 3-D printed collimator filled with aqueous ultrasound gel at an incidence angle of 40°(Tufail et al., 2011). A 32-channel electrode array is inserted using a stereotaxic arm into the targeted brain area (i.e. primary somatosensory cortex S1, ML: −3 mm, AP: −0.84 mm, depth: 1 mm, Fig. 3A) prepared through craniotomy, and histologically confirmed through a coronal brain slice post hoc. The burr hole through the skull with an approximate diameter of 2 mm is shown in a sagittal CT image (Fig. 3B). The stimulation dynamics of the tFUS waveforms consist of tone-burst sinusoidal waves with constant UFF (500 kHz), and TBD (200 μs), and varied UPRF, spanning five levels between 30 – 4,500 Hz (Fig. 2B, Table 1 in STAR Methods). The duration of inter-sonication interval (ISoI) is 2.5 seconds per trial. A hydrophone-based 3-D ultrasound pressure scanning system was used to experimentally obtain measurements of the tFUS spatial profile behind a freshly-excised rat cranium. To illustrate the tFUS spatial distribution, the spatial profile of the ultrasound in the x-y plane is superimposed on a rat cranium (Fig. 2C). The ultrasound spatial map from the coronal view (y-z plane) is reconstructed, in which mechanical energy is distributed along a coronal beam up to a depth of 4 mm, but spatial peak energy is located within 1 mm from behind the skull (Fig. 2D). This illustrates, the 40-degree angled incidence (Tufail et al., 2011) leads to a shallowed targeting at the rat cortex, dissipating the majority of ultrasound energy through the skull. Angled tFUS stimulation is the preferred method for studying the cortex, as its activation pattern is shallower than normally incident tFUS. Numerical simulations using a pseudo-spectral time domain method of the spatial distributions of the ultrasound pressure throughout the imaged rat skull were conducted in order to account for potential acoustic reflections from the skull base in the small enclosed cavity (Fig. 2E-F).

**Table 1.** Administered tFUS Conditions with Featured Parameters

| tFUS Conditions* | UPRF (Hz) | UDC (%) |
| --- | --- | --- |
| UPRFx1 | 30 | 0.6 |
| UPRFx10 | 300 | 6 |
| UPRFx50 | 1,500 | 30 |
| UPRFx100 | 3,000 | 60 |
| UPRFx150 | 4,500 | 90 |

\* Except for the ultrasound pulse repetition frequency (UPRF) and thus the ultrasound duty cycle (UDC), all of the listed tFUS conditions use the same UFF, UD, ISol, TBD, and spatial-peak temporal-peak intensity $I_{sptp}$ .

**Figure 2.**
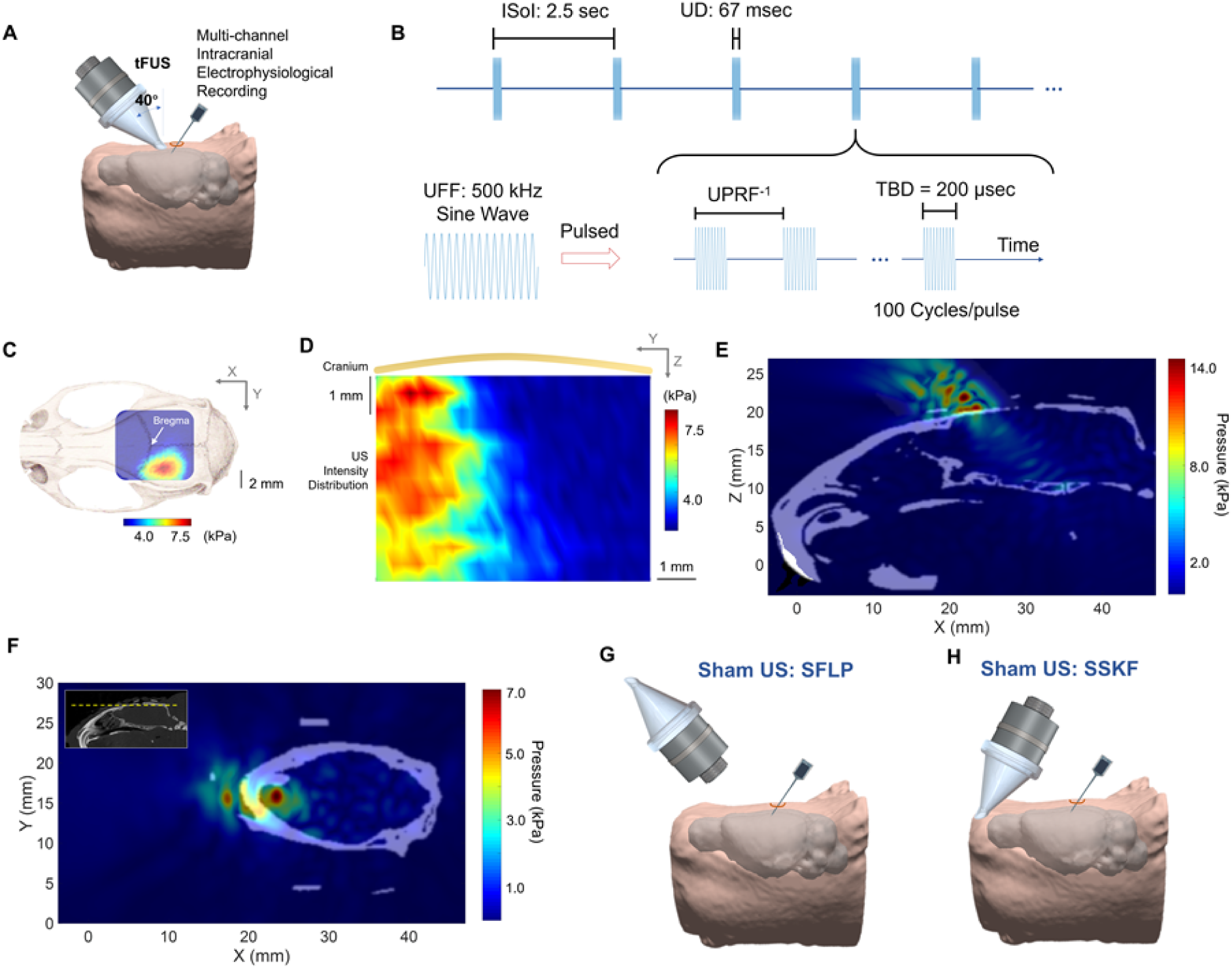
*In Vivo* Ultrasound Stimulation at Somatosensory Cortex. **(A)** Collimator guiding focused ultrasound (US) to hair-removed scalp of an anesthetized rat model with an incidence angle of 40°, and a 32-channel electrode array inserted at an incidence angle of 40° into the left primary somatosensory cortex (S1) prepared through a craniotomy. The rat head and brain model were 3D reconstructed from T_2_-weighted (T2W) MRI images (Valdes-Hernandez et al., 2011). (**B**) The temporal profile of tFUS. 100 cycles of sinusoidal wave formed a single ultrasound pulse, which generated a tone-burst at an ultrasound fundamental frequency (UFF) of 500 kHz for a tone burst duration (TBD) of 200 μs. Such ultrasound pulses were repeated at certain UPRF for a corresponding number within the ultrasound duration (UD) of 67 ms. The inter-sonication interval (ISoI) was 2.5 seconds. (**C-D**) One transverse (X-Y plane) and one coronal (Y-Z plane) scans of ultrasound pressure distribution under the cranium using a hydrophone-based US field mapping system. After transmission through the *ex-vivo* skull top, spatial-peak ultrasound pressure is measured at 7.9 kPa, spatial-peak temporal-average intensity (I_spta_) is 15.2 mW/cm^2^ at a UPRF of 1,500 Hz, and spatial-peak temporal-peak intensity (I_sptp_) is 405 mW/cm^2^. (**E**) A 3D computer simulation investigating the quantitative maximum pressure distribution at one sagittal (X-Z) plane inside the rat skull (in white and gray colors) when the tFUS is directed to the S1 with an incidence angle of 40°. (**F**) A transverse (X-Y plane) view of the ultrasound pressure field. The yellow dashed line shown in the inset indicates the location of the presented transverse plane. (**G**) The experimental setup of the sham US condition as a negative control (SFLP). The ultrasound transducer was flipped its aperture by 180°, although the pulsed ultrasound and the intracranial recordings were maintained. (**H**) The experimental setup of another sham US condition as a control for bone-conduction (SSKF). The ultrasound incidence took place at an anterior location of the skull, and the aperture was kept transmitting ultrasound. Another simulation study, as shown in Fig. S1A, mapped the ultrasound pressure (in red color) at the sagittal (X-Z) plane when the tFUS transmitted to the control location, SSKF. The quantitative map is shown in Fig. S1B.

**Figure 3.**
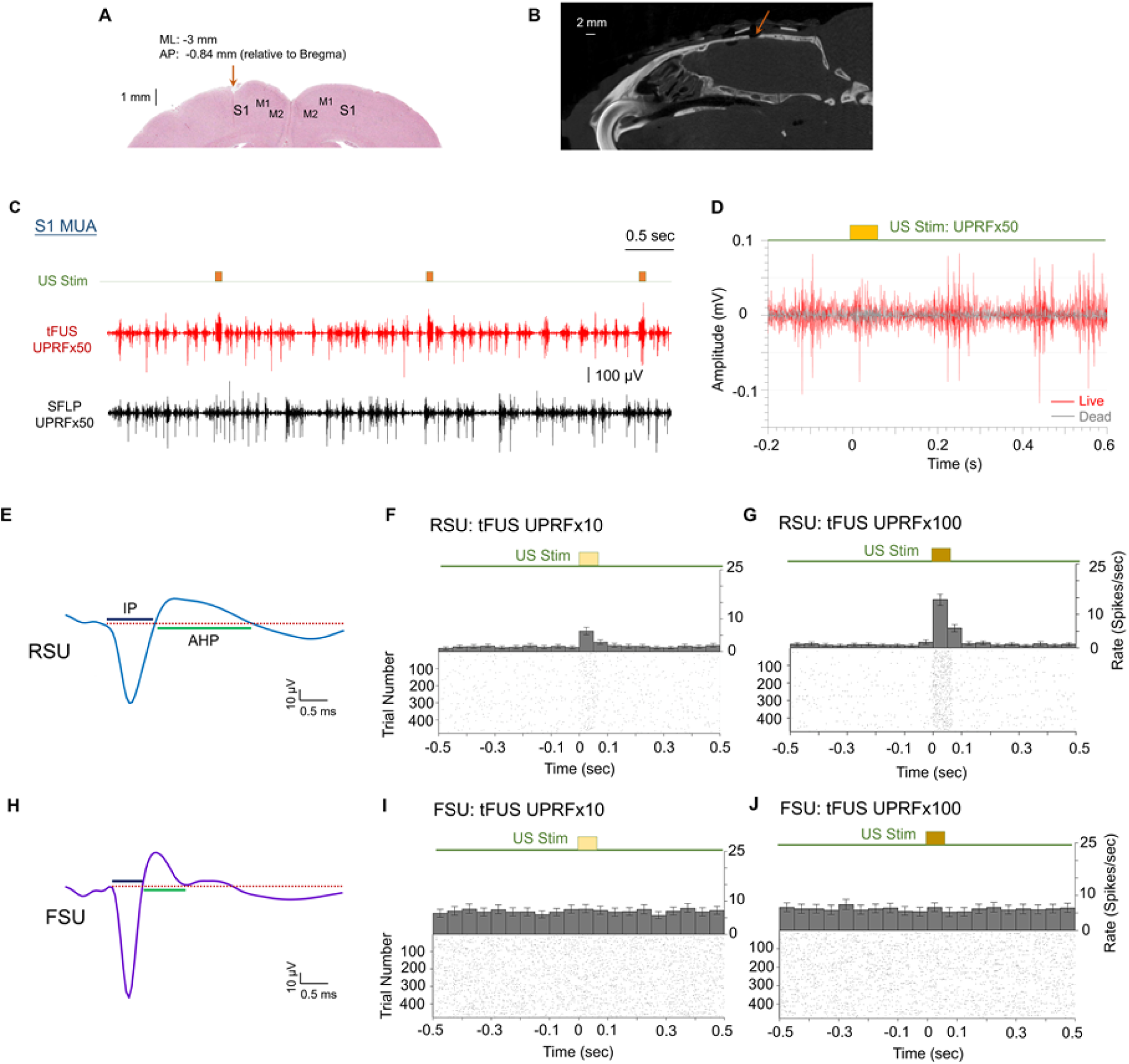
*In Vivo* Electrophysiological Recordings. (**A**) The spatial coordinates of electrophysiological recordings. The coronal brain slice shows the location of electrode insertion as a physical breakage with the insertion depth of 1 mm at the left S1. ML denotes the medial lateral distance from midline; AP denotes the anterior posterior distance from Bregma. (**B**) A sagittal view of the Micro-CT image captured the surgical burr hole (approximate diameter: 2 mm) on the top of cranium. (**C**) An example of acquired multi-unit activity (MUA) from the S1 using tFUS with UPRF=1500 Hz. The timing between the ultrasound-induced action potentials and the administered stimulations are exemplified by 4 trials. The sham condition, SFLP shows a silence of such time-locked MUA. (**D**) Wavelet denoised multi-unit activity waveforms of recorded stimulation response in live (red trace) and dead (gray trace) animal under tFUS stimulation at UPRF=1500 Hz to demonstrate the substantial removal of stimulation artifacts. (**E**) A typical example of an RSU separated from the recorded MUA. The waveform features, i.e. time durations of initial phase (IP) and afterhyperpolarization (AHP) were employed to conduct the units’ separation. (**F-G**) The peri-stimulus time histograms (PSTHs, bin size: 50 ms) and raster plots of the spiking unit (IP duration mean: 850 μs; AHP duration mean: 1850 μs) responding to the UPRF of 300 Hz (F) and 3000 Hz (G). (**H**) An example of an FSU identified with short AHP duration (H). (**I-J**) The PSTHs and spiking raster plots of such a spiking unit (IP duration mean: 700 μs; AHP duration mean: 600 μs) in response to the two tFUS conditions with UPRF of 300 Hz (I) and 3000 Hz (J). The time histograms and return plots of the inter-spike interval (ISpI) are depicted in Fig. S3B-C. All of the temporal dynamics are computed across 478 trials, with each trial lasting 2.5 s. The applied ultrasound conditions are also described in Table 1 (see STAR Methods). Data are shown as the mean±95% confidence interval in the PSTHs.

To control for possible confounds due to acoustic and electromagnetic noise in the experimental setup, two sham US conditions were conducted. The sham condition, SFLP transmits ultrasound waves in air, directed 180 degrees away from the skull (Fig. 2G). Another sham condition, SSKF, in which tFUS was directed at a secondary control site physically away from the target site, is used to control for the skull bone conduction (Fig. 2H). Numerical stimulations were also conducted for this SSKF sham experiment (Fig. S1A-B). The spatial specificity of the ultrasound-induced brain activities was also demonstrated in the supplements (Fig. S2).

During tFUS stimulation, increased time-locked neuronal firing is observed in recorded MUA as compared to SFLP sham conditions (Fig. 3C). Furthermore, to eliminate the possibility of tFUS producing mechanical deformation related potentials on the recording electrode, a comparison of the recorded signal is conducted in the same animal before and after euthanasia. Fig. 3D illustrates that no significant tFUS related electrical artifacts are observed on the recording electrode in the dead animal. With necessary preprocessing (see STAR Methods), all tFUS aligned activations observed on the electrode are deemed to be the neural responses to the ultrasound depositions.

### Local Activations are Preserved in tFUS Setup When Auditory Pathway is Blocked

To test whether the induced neural activations is locally directed by the tFUS, we perform control studies in chemically deafened rats under tFUS stimulation. The study methods are depicted as in Fig. 4A. The auditory brainstem response (ABR) tests were conducted through subcutaneous needle electrodes to measure the rats’ percepts to an auditory stimulus, and were conducted before and 72-hour after chemical injections of furosemide and gentamicin. Fig. 4B shows the results from the ABR test of a naïve rat and Fig. 4C shows a complete lack of response to auditory stimuli 3 days after the chemical injections. The deafening is conserved across different frequency ranges of auditory stimuli, an indication of widespread hearing loss.

**Figure 4.**
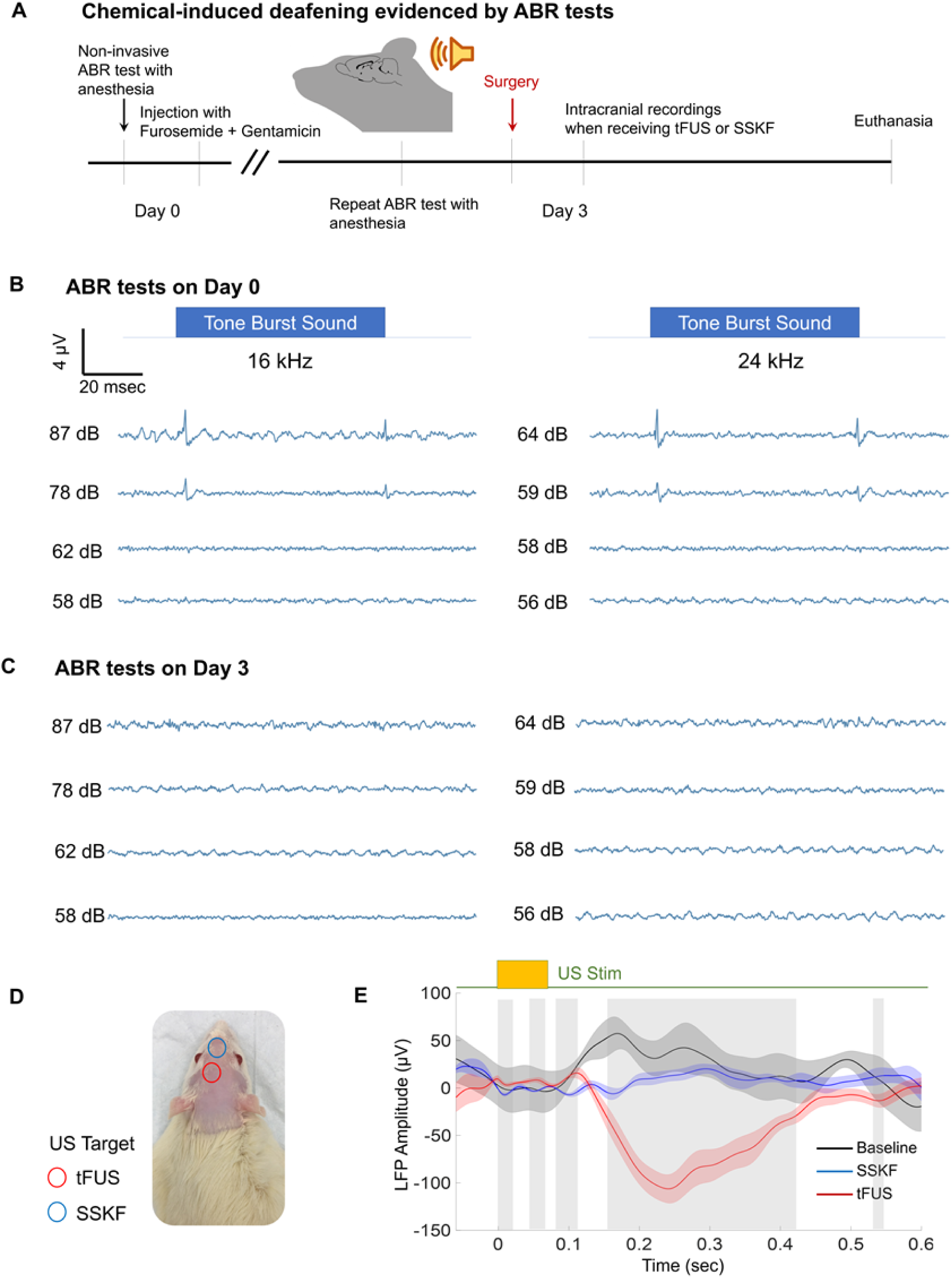
tFUS Induced Local Brain Activity on A Chemically-induced Deafened Rat. (**A**) The experimental protocol for creating chemical-induced deafening model. The deafening will be tested by pre- and post-chemical injection auditory brainstem responses (ABR) tests on Day 0 and Day 3, respectively. A combination of injected chemicals includes furosemide and gentamicin. (**B-C**) ABR test results before and after chemical injection. Four levels of sound intensity with 16 and 24 kHz center frequencies were used. On Day 3, post-injection ABR test results show significantly reduced auditory brainstem responses to the levels of sound intensities. (**D**) On a photo representation of the rat in anesthesia after chemically deafened, ultrasound at the targeting location marked as red circle, or at the control location identified as blue. (**E**) LFP recorded at the S1 area and averaged across 400 trials, and presented with the mean value (solid line) and standard error of the mean (shaded areas). The employed ultrasound condition is UPRFx50. Comparing with results from the sham condition, i.e. SSKF, significant differences with multiple comparison correction were observed and marked with vertical gray bars (*p* < 0.005).

After deafening is ascertained, tFUS was applied at S1, similar to experiments on healthy rats. Fig. 4E illustrates the local field potentials measured at S1 during baseline, tFUS stimulation and SSKF sham condition (UPRF = 1500 Hz, I_spta_ = 15.2 mW/cm^2^). Across 400 trials, time aligned activation is observed in the LFP during tFUS, originating at approximately 100 ms after tFUS onset. The standard error of mean potentials are illustrated; gray shaded regions show statistically significant differences amongst the average LFP traces, and note that the significant different temporal regions are largely in the time segment of 155-417 ms after sonication, which is outside the timescale of auditory induced propagations observed in LFP data by (Guo et al., 2018). This activation is significantly different compared to the sham condition (Fig. 4D) at an anterior skull location tested through permutation-based non-parametric statistics with false discovery rate (FDR) multiple-comparison correction (*p* < 0.005).

### Rat Somatosensory Cortex Exhibits Cell-type Specific Response to tFUS

All recorded action potentials from the 32-channel electrode array were sorted based on the spike waveforms and inter-spike intervals (ISpI). Regular-spiking (presumably excitatory) (RSU) and fast-spiking (presumably inhibitory) units (FSU) are thus identified based on the temporal dynamics of the action potential waveform, shown in Fig. 5A, D and F (Mountcastle et al., 1969; Murray and Keller, 2011; Simons, 1978). The features extracted are the durations of initial phase (IP) of the action potential, i.e. from onset to the re-crossing of baseline, and afterhyperpolarization period (AHP), i.e. from the end of the IP to its re-crossing of baseline, shown in Fig. 3E. The differences in these features have been associated with differences of ion channel types and distributions in the neuronal cell membrane. We thus hypothesize that the RSU and FSU will have distinct responses to various tFUS stimulation sequences due to their intrinsic cellular differences, which is to be statistically tested given one of the putative mechanisms of ultrasound neuromodulation is that mechano-sensitive ion channels mediate the acoustic induced neural effects (Tyler, 2011).

**Figure 5.**
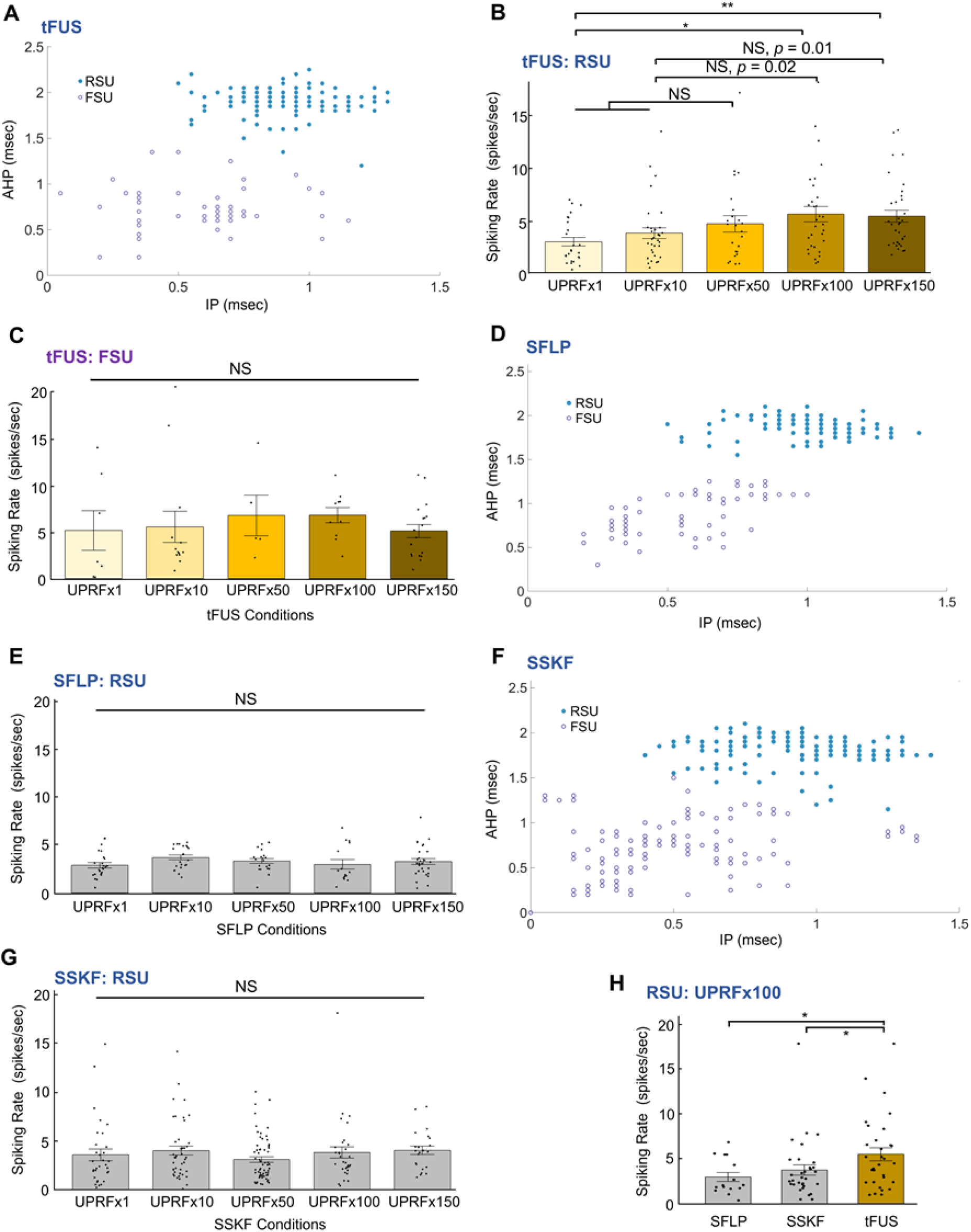
Cell-type Selective Responses to tFUS and Sham US Conditions. (**A**) k-means cluster analysis of 199 single units identified from 6 rats with blue solid circles depicting the RSUs while purple circles representing the FSUs. The majority is classified as the RSU with longer IP and/or AHP durations than the FSU. These spiking units are recorded and identified under the influence of the administered anesthesia (xylazine) and analgesic drugs (ketamine), in which the durations of both IP and AHP are observed to be longer than the results reported in literature (Bruno and Simons, 2002; Murray and Keller, 2011) for different sedative approaches. (**B**) The 146 RSUs significantly differ their responses to different tFUS conditions. Data are shown as the mean±s.e.m., with statistical comparisons made through Kruskal-Wallis one-way ANOVA on ranks and *post hoc* one-tail two-sample Wilcoxon tests with Bonferroni correction for multiple comparisons. \**p* < 0.005, \*\**p* < 0.001. NS, not significant. s.e.m., standard error of mean. (**C**) The FSU group, which consisted of fewer neurons (N = 53, marked as purple hollow circles), was observed to have a higher mean spike rate than RSUs, whereas this FSU group showed no significant effect by different UPRF levels. Data are shown as the mean±s.e.m., statistics by Kruskal-Wallis H test. (**D**) In one of the sham conditions, i.e. SFLP, the cluster analysis for another identified 179 single units. 110 RSUs and 69 FSUs were separated. (**E**) RSUs showed a homogeneous lack of response to the SFLP sham ultrasound conditions. No significant effect by the ultrasound conditions was observed through statistics by Kruskal-Wallis H test. Data are shown as the mean±s.e.m. NS, not significant. s.e.m., standard error of mean. (**F**) In another sham control, i.e. SSKF, for investigating any potential effects of bone-conduction, the cluster analysis for another identified 311 single units. 192 RSUs and 119 FSUs were separated. (**G**) RSUs also showed a homogeneous lack of response to the SSKF sham ultrasound conditions. No significant effect by the ultrasound conditions was observed through statistics by Kruskal-Wallis H test. NS, not significant. s.e.m., standard error of mean. (**H**) Group comparisons between SFLP and tFUS, and between SSKF and tFUS of RSUs’ spiking rates responding to the UPRFx100 level. Data are shown as the mean±s.e.m., statistics by two-tail Wilcoxon test. \**p* < 0.05.

After recording the MUAs from the first group of wild-type male Wistar rats (N = 9), we studied the neural effects of the administered pulsed tFUS through intracranial MUA recordings. Using peri-stimulus time histograms (PSTH), we found a significant increase of spike rate (6.23±1.10 spikes/sec, Fig. 3F) in a possible regular-spiking somatosensory cortical neuron (mean spike waveform IP: 0.85 ms, AHP: 1.8 ms) when stimulated with a tFUS condition (UPRF=300 Hz, I_spta_=3.0 mW/cm^2^), and observed a further increased spiking rate (14.35±1.65 spikes/sec, Fig. 3G) in response to the increase sonication (UPRF=3000 Hz, I_spta_=30.4 mW/cm^2^). For a more intuitive comparison, increases of spiking as a function of time along 478 consecutive trials is demonstrated with the raster plot (Fig. 3F-G), in which the density of spiking events increases during the ultrasound stimulation. For another identified RSU, the comparisons are shown between the tFUS and sham conditions. Using the UPRF of 1500 Hz, a significant bursting peak, a type of non-Poissonian, tFUS-mediated behavior is shown in the time histogram and the return plot of ISpI (Fig. S4A, B). Unsurprisingly, the SFLP and SSKF conditions do not exhibit such distinct spiking increase due to stimulation seen from either PSTH, raster (Figs. S4C, E) and return plots (Fig. S4D, F).

In contrast, a fast-spiking cortical neuron (Fig. 3H) with shorter durations of IP (mean: 0.7 ms) and AHP (mean: 0.65 ms) shows a more homogeneous PSTH distribution in response to the levels of tFUS treatment (e.g. UPRFx10 and UPRFx100 in Fig. 3I-J) tested using the ultrasound setup shown in Fig. 2A. The return plots (Fig. S3B-C) illustrate a fast spiking behavior with a shorter refractory period than the RSU spiking unit identified in Fig. 3E. As seen in the example (Fig. 3I), FSU firing rates are not disturbed by tFUS, i.e. that the firing rate (7.6±1.2 spikes/sec) is not significantly altered by the US stimulation (UPRF=300 Hz, I_spta_=3.0 mW/cm^2^) comparing to pre-stimulus rates (e.g. 7.5±1.2 spikes/sec at the bin of [-0.05, 0] s). Interestingly, for FSUs, we observed no significant changes in spike rates (6.5±1.3 spikes/sec) even when ultrasound was administered at a UPRF 10 times higher (UPRF=3000 Hz, I_spta_=30.4 mW/cm^2^, Fig. 3J). Furthermore, the return plots (Fig. S3B-C) indicate that although this fast-spiking neuron does not significantly change its rate of firing action potentials in response to tFUS, there is a possible trend of changing the spiking pattern of its bursting mode.

Given these case studies, we pursued a statistical investigation of the behavior of different neuron types in response to tFUS and sham conditions, across multiple levels of UPRFs. In the tFUS group, we compiled all single unit activities during the UD of 67 ms recorded from the rats and separated them into RSUs and FSUs using k-means cluster analysis (Fig. 5A). 199 identified single units were separated into the two groups, with the RSU group containing 146 units. The sample sizes are unbalanced due to the prevalence of each cell types in the cortex. The baseline firing rate of each single unit in different ultrasound sessions, which may be affected by anesthetic depth, is controlled by randomizing the order of tFUS conditions.

In the RSU group, we found UPRF levels have a statistically significant effect on the neuron firing rates (Kruskal-Wallis chi-squared =14.45, p = 0.006), indicating that the RSUs respond to the different tFUS conditions with different firing rates (Fig. 5B). More specifically, the spike rate increases have shown a positive correlation with increasing UPRFs. Through the multi-comparisons, we found that in an anesthetized rat model, the RSUs significantly increase their firing rates in response to UPRFs at 3000 and 4500 Hz when both comparing to that induced by a low UPRF at 30 Hz (Fig. 5B, UPRFx1 vs. UPRFx100: p = 0.003; UPRFx1 vs. UPRFx150: p = 0.0004). Whereas in the FSU group (Fig. 5C), no significant difference between tFUS conditions could be found (Kruskal-Wallis chi-squared = 4.34, p > 0.3). This implies that the spike rates of the FSUs were not significantly altered by the levels of UPRF, consistent with the case study (Fig. 3I-J and Fig. S3A).

The contrast between the responses observed in these two different neuron types suggested a cell-type selective mechanism by tFUS. Unsurprisingly, the RSU group did not show significant differences among the five levels of sham US conditions, including the SFLP (Kruskal-Wallis chi-squared = 6.58, p > 0.15, Fig. 5E) and SSKF (Kruskal-Wallis chi-squared = 6.32, p > 0.15, Fig. 5G) shams. Moreover, as seen in Fig. 5B, the first significant difference shows up at the UPRF of 3000 Hz; when the tFUS was compared with the corresponding sham conditions, significant increase of spiking rates were found in the tFUS against SFLP and SSKF at this UPRF (Fig. 5H, tFUS vs. SFLP: p = 0.033; tFUS vs. SSKF: p = 0.048).

In summary, this is the first time that the intrinsic, distinct difference in responses of RSUs and FSUs to the change of UPRF is found. Since the length of the refractory period determines the minimum time between neuronal firings, it is not surprising to see that FSU spikes faster than the RSU (Fig. 3F-G vs. Fig. 3I-J). However, the spike rate responses to tFUS between these two groups is a new finding (Fig. 5B-C). When administered with a low UPRF (i.e. 30 Hz), the RSUs do not respond significantly to tFUS stimulation comparing to the shams (Fig. 5B, E and G), meanwhile FSUs also maintain a stable spiking state during the sonication (Fig. 3I-J, Fig. 5C and Fig. S3A). The observed responses suggest cortical neurons with different action potential shapes, hence different distribution of ion channels, in terms of ion channel types or relative quantity, have distinct response patterns to tFUS UPRF or duty cycle tabulated in Table 1 (see STAR Methods). While FSUs are maintaining the activities across all UPRF frequencies, RSUs only exhibit increased firing rate during high UPRFs. This could explain why when stimulating at low UPRFs, some previously reported studies have observed tFUS suppression effects (Legon et al., 2018; Legon et al., 2014). The observed effects may not simply be due to the lack of facilitation in the neural circuit, but rather the leading inhibition in the network could play a significant role.

To further assess our cell-type specificity hypothesis, we tested our results in transgenic mouse models with PV (N = 53, from 3 mice) and CamKIIa (N = 50, from 2 mice) cortical neurons identified by response to optical stimulations. Fig. S7A illustrates the *in vivo* experimental setup combining optical stimulation, tFUS and multi-channel intracranial electrophysiological recordings. The optogenetic stimulation (wavelength = 465 nm) locally activates a subpopulation of channelrhodopsin expressing neurons, based on the MUA recording from our optrode, we can thus identify the cell type of our recording neurons (Fig. S7A). The waveforms of action potentials of each neuron are illustrated together with the spike rates when receiving the optical stimuli (Fig. S7B). Since the activation of channelrhodopsin has been linked to changes in neuronal baseline spiking dynamics, all optogenetic stimulations are administrated after completing all tFUS recording sessions. Between PV and CamKIIa neurons, we observed significant differences in the action potential waveforms regarding the IP and AHP phase durations, validating our method for separating neuron population in rats (Fig. S7C, *p* < 0.05). In PV neurons, spike rates are higher under tFUS stimulation at UPRF of 30 Hz (Fig. S7D, *p* < 0.001), while in CamkIIa neurons, significantly higher spike rates are observed during tFUS stimulation at UPRF of 300 Hz (Fig. S7E, *p* < 0.01). Such differences in response to tFUS UPRFs across these two neuron types show further evidences of cell-type specific effects of tFUS.

### tFUS Induces Long Term Changes in Synaptic Connectivity

In order to further examine the cell type selectivity of tFUS, we set out to explore the effects of tFUS UPRF on activating networks in the deep brain (Fig. 6A). In this study, we examine whether it is plausible to use tFUS to induce long term synaptic plasticity in the hippocampus, and whether the frequency of tFUS UPRF has a significant effect on outcome. For a deeper brain target, we designed a third collimator (see Methods) to allow normal incidence of tFUS at the scalp. We thus see a much higher and deeper spatial-peak ultrasound pressure (Fig. 6C-D) field than that when using the angled incidence (Fig. 2C-F). A local maximum ultrasound pressure can be observed at a depth of 3.5 mm in the hydrophone scanning (Fig. 6D), and computer simulations of the pressure field (Fig. 6E-F) demonstrate that ultrasound energy is able to penetrate deep and aim at the subcortical regions with the normal ultrasound incidence.

**Figure 6.**
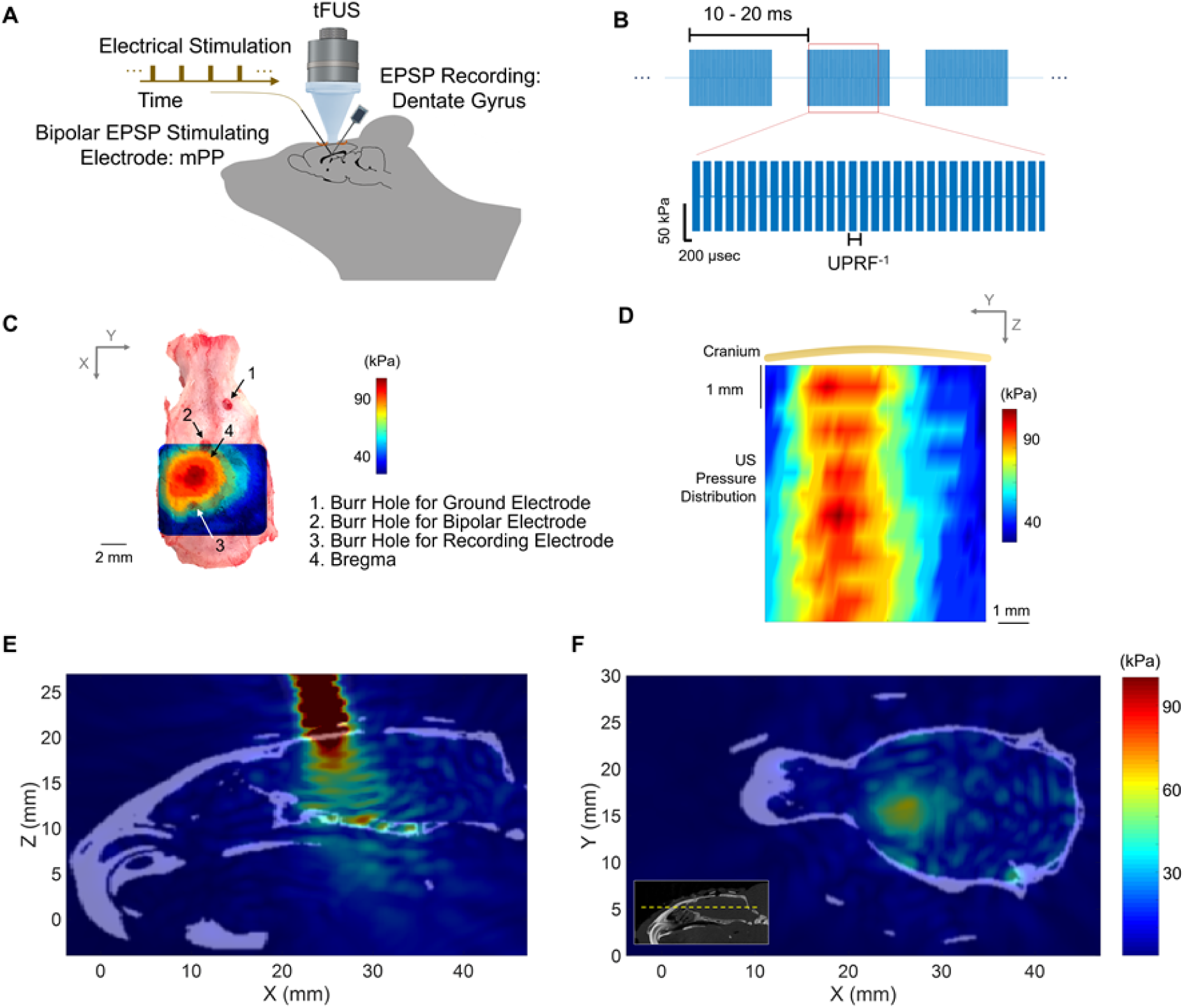
*In Vivo* Ultrasound Stimulation at Deep Brain. (**A**) Schematic of experimental setup. tFUS is delivered with normal incidence to the skull. fEPSP stimulating electrode inserted at 40 degrees and recording electrode inserted at 50 degrees incidence angle. (**B**) tFUS stimulation waveform during long-term effect induction. Trains of pulsed tFUS at various UPRFs (3 - 10 kHz) are delivered at 50 Hz (20 ms) intervals at 60% duty cycle to mimic the frequency of long-term potentiation (LTP) induction using electrical stimulation. (**C-D**) Ultrasound field mapping of one transverse (X-Y plane, C) and one coronal (Y-Z plane, D) scans of ultrasound pressure distribution during tFUS induction under the cranium using a hydrophone-based US field mapping system. (**E-F**) A computer simulation maps the ultrasound pressure field inside the skull cavity. One sagittal (X-Z plane) view is illustrated and co-registered with the skull CT image (E); One transverse (X-Y plane) view (F) with its location indicated by the yellow dashed line in the inset of (F) is shown for depicting the ultrasound pressure field at the deep brain.

To test whether tFUS can induce frequency encoded potentiation in the synapse, we attempted the induction of LTP using pulsed tFUS in the naïve rats (N = 9). The main goal of this set of experiments is to investigate the ability to elicit long-term changes in synaptic strengths using tFUS. We applied pulsed tFUS stimulation with various UPRFs at 50-100 Hz sonication frequency (Fig. 6B) in order to emulate the effects of high frequency electrical stimulation of LTP in the dentate gyrus. Field response is assessed from the maximum descending slope of the fEPSP, shown in Fig. 7B and D. Given that when the UPRF reaches 3 kHz, the cortical RSUs exhibit significant increased activities (Fig. 5B and H), we set the UPRF starting from 3 kHz for studying the transcranial neural effects at the deep brain. In the nine rats, instead of LTP, LTD was observed in the fEPSP when tFUS stimulation was applied with UPRF of 3-10 kHz (Fig. 7E). LTD was observed to persist 30 minutes after stimulation cessation (Fig. 7A). As seen in Fig. 7A, the fEPSP slope significantly decreases after tFUS stimulation and returns toward baseline over time. After all tFUS experiments, LTD is elicited using low frequency (1 Hz) electrical tetanus stimulation to validate the correct localization of neural pathway (Fig. S8). Sham experiments with tFUS delivered at 180 degrees away from the skull demonstrate that LTD does not occur without the presence of tFUS (Fig. 7C-D). Averaged fEPSP in Fig. 7B and D show changes in fEPSP immediately pre or post tFUS stimulation, averaged across 5 minutes. The slope of the descending segment of fEPSP after the electrical pulse at 0 ms is used to calculate fEPSP slope.

**Figure 7.**
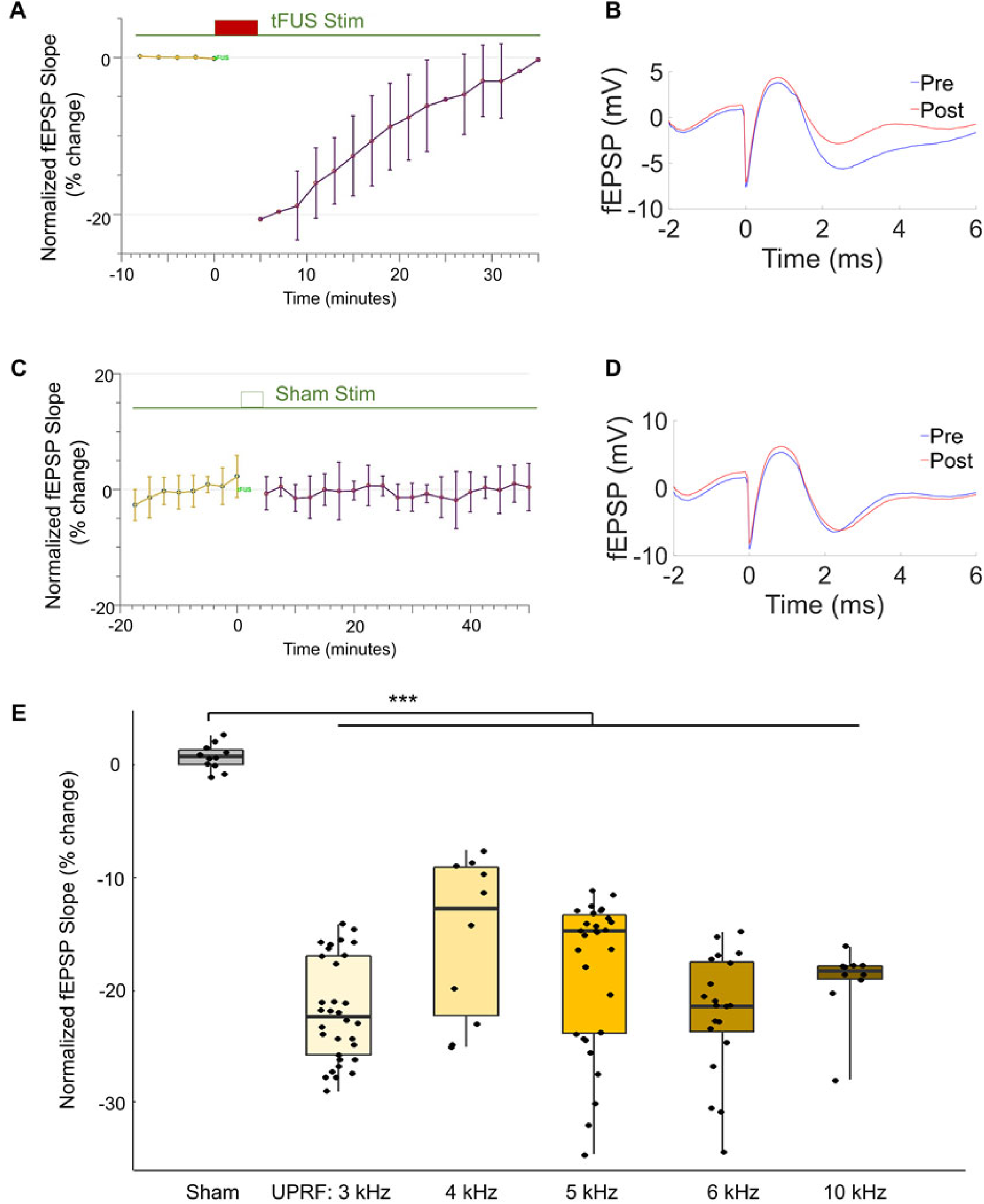
Induction of Long Term Depression (LTD) in Rats. (**A-B**) LTD induced by pulsed tFUS at different UPRF from 3 to 10 kHz. Stimulation at the medial performant path recording from DG (N = 9). The red block denotes the timing of administering tFUS for 5 minutes. The waveforms of fEPSP recorded in tFUS stimulations (B). The blue line represents the fEPSP before the stimulation, and the red line indicates the one after the stimulation. Data shown in (A) as mean±s.t.d.. s.t.d. standard deviation. (**C-D**) The sham tFUS was delivered by flipping ultrasound transducer 180 degrees away from skull. All other procedures held constant. Average fEPSP waveforms pre- and post-stimulation are shown in (D). Data shown in (C) as mean±s.t.d.. s.t.d. standard deviation. (**E**) Data are shown in the boxplot as the median with 25% and 75% quantiles (lower and upper hinges). statistics by one-way ANOVA test with post hoc Tukey’s Honestly Significant Difference Procedure. \**p* < 0.05.

Based on our hypothesis, we expected to observe LTP after tFUS stimulation since tFUS was applied at the same frequency as the high frequency tetanic stimulation used throughout LTP literature. The observed results did not show LTP, suggesting that the temporal encoding using tFUS does not share the same efficiency and/or mechanism as electrical tetanus stimulation. However, the demonstrated long-term effect is a promising new feature of tFUS stimulation to be employed as a potential non-invasive therapeutic neuromodulation technique. Our results suggest that tFUS can be used to encode time dependent stimulation paradigms into neural networks and non-invasively elicit long-term changes in the strength of synaptic connections.

In order to investigate whether tFUS UPRF has an effect on strength of LTD induction, we examined a range of UPRFs from 3 to 10 kHz. Shown in Fig. 7E, all of the UPRFs applied are significantly different when compared to sham stimulation, based on one-way ANOVA tests with a post hoc Tukey’s honestly significant difference test procedure to adjust for difference in sample size. Although there are no observed statistically significant differences between the sample groups, overall, we observed LTD across all sample groups. This phenomenon suggests tFUS UPRF is not correlated with LTD, however the strength of LTD induction may be affected by tFUS UPRFs.

## DISCUSSION

In the present study, we set out to use multi-channel intracranial recordings to test our hypotheses of tFUS’s ability to induce cell-type specific stimulation and induce long term changes in synaptic connectivity for the first time. Based on the results reported above, we have gained better understanding of the tFUS parameter space, and thus can furthermore infer on the mechanism of action of tFUS stimulation. Besides macroscopic perspectives reported in literature, uncovering the underlying neuronal mechanism requires a detailed inspection of how neurons respond to a vast set of acoustic parameters. Recordings using multi-channel intracranial electrophysiology allows us to examine the neuron cellular dynamics with high spatial and temporal specificity.

### Controls for Confounding Effects

Based on a simultaneous EEG source imaging technique applied on rats reported previously (Yu et al., 2016), observations at the auditory cortex can be induced as a secondary activation when tFUS is not directed at the auditory cortex (Niu et al., 2018). Therefore, in order to examine the auditory side effects (Guo et al., 2018; Sato et al., 2018) of the tFUS setup, presumably due to tFUS induced mechanical vibrations transmitted through the skull, we conducted tFUS stimulation after inducing permanent chemical deafening on rats (Fig. 4A). Our observations of a significant rise (Fig. 4D-E) in local field potential (LFP) at the S1 cortical region on the subject with significant reduction in hearing threshold level (Fig. 4B-C), suggest that side-effect activations in the auditory cortex from tFUS induced hearing percepts do not dictate activation of S1 cortex. Unsurprisingly, in this deafened model, the tFUS-induced LFP resembles the reported LFP waveform by Tufail et al. (Tufail et al., 2010) on the mice model. Similar ultrasound parameters (e.g. UFF and UPRF) were employed in both studies. In spite that the presence of the auditory pathways is disabled in the neural networks, significant tFUS activations can still be observed, providing direct evidence that our tFUS setup is able to induce local activations which are independent to the auditory perceptions.

Although the chemicals used in this group of experiments have been shown to damage auditory pathways, we do not have confirmation of whether these drugs present toxicity to other parts of the neural network. Therefore, one should be cautious in critically evaluating recordings gathered from these deafened rats. In order to control for auditory confounding effects leading to false positives in our tFUS stimulation on healthy rats, we performed two different sham conditions along with tFUS stimulation.

Furthermore, we provide a possible explanation to the previously observed auditory confounding effects (Guo et al., 2018; Sato et al., 2018), whose observations on the small rodents might also be fundamentally confounded by the extensive ultrasound field within the rodent skulls. We speculate auditory activations may be due to ultrasonic standing waves in the skull cavity resultant from reflections from the base of the skull. Such wave pattern appears to be more extensive when using even lower UFF (Guo et al., 2018) or even smaller object dimensions of a mouse head (Sato et al., 2018). Conventional *ex-vivo* pressure mapping does not easily capture these standing waves due to the negligence of the major reflections at the skull base. Simulation results illustrate the estimated distribution of these standing waves (Fig. S1C-J), including the opened-skull simulations (Fig. S1G-H) on 1.5 times enlarged head size to compensate the potential size differences between the Wistar rat and guinea pig models (Guo et al., 2018), although both models have a similar skull size (Albuquerque et al., 2009). In the small skull cavities, the ultrasound pressures are concentrated at the local target region as expected (Fig. 2E-F, Fig. 6E-F), however lower amplitude standing waves are still observed to be an extensive existence throughout the brain, including the auditory cortex especially when a lower UFFs of 220 kHz and 350 kHz were employed as in Fig. S1E-H (Guo et al., 2018) and Fig. S1I-J (Kim et al., 2014), respectively.

Since our stimulation location is at the primary somatosensory cortex, control studies were conducted to examine whether activations recorded in the S1 could be due to somatosensation rather than direct activation of neurons in the S1. In a study examining the effect of tFUS on anesthesia recovery time, described in supplementary materials (Fig. S6), rats with tFUS stimulation directly at S1 (UPRF=1500 Hz) recovered from anesthesia significantly faster than rats during sham US. Controls were tested for rats for auditory percepts coupled and not coupled to the skull, and rats with peripheral stimulation at the contralateral hind limb to control for somatosensation. Control rats with auditory percepts coupled to the skull has tFUS focused on the anterior skull location, away from S1; control rats with auditory percepts not coupled to the skull have ultrasound directed 180 degrees away from S1. This study suggests that our tFUS experimental setup can elicit direct stimulation of the rat brain without dependence on confounding effects of auditory percepts and somatosensation.

### The Selectivity between Excitatory and Inhibitory Neurons by Tuning The UPRF-related Duty Cycle

The results reported above are *in vivo* evidences that subsets of neurons, grouped by their action potential waveforms, respond differently to tFUS stimulation UPRFs. This provides insight into results previously reported by other groups studying tFUS (King et al., 2013; Lee et al., 2016b; Legon et al., 2014; Tufail et al., 2011). Our results suggest that tFUS interacts with neurons based on ion channel dynamics or neuronal morphology. Hence, the intrinsic differences in ion channel dynamics between different neuron types or the distinct profiles of dendritic arbors contribute to differences in the neuron’s response to tFUS.

The ultimate goal of studying the mechanisms of tFUS is to translate the technology to clinical utility. Previous studies have shown that inhibitory effect of tFUS was found at the primary somatosensory cortex (Legon et al., 2014) and thalamus (Legon et al., 2018) in humans, in which the same ultrasound parameters were employed (single-element transducer with UFF = 500 kHz, UPRF= 1 kHz, UD = 500 ms, UDC = 36%). In contrast, other studies on humans have shown excitatory effects that the primary visual cortex was directly excited with a specific ultrasound administration (UFF = 270 kHz, UPRF = 0.5 kHz, UD = 300 ms, UDC = 50%) (Lee et al., 2016b), and simultaneous stimulating capability of tFUS were also shown at primary and secondary somatosensory cortices (UFF = 210 kHz, UD = 500 ms) (Lee et al., 2016a).

Given these parameter-dependent studies, the effects of tFUS achieved by these two groups are sometimes observed to be inconsistent or even contradictory on healthy, awake subjects. We cannot conclude from these studies whether the observed behavior was due to overall activation or suppression of neural activity due to changes in tFUS parameters, or if the ultimate behavior was due to selective modulation of the neural network. The modified NICE model was proposed to unify the ultrasound parametric space and predict either excitatory or inhibitory effects at a neuronal level (Plaksin et al., 2016). As we set out to explore the tFUS parameter space in the *in vivo* brains, we set our administered duty cycle of tFUS to the five levels (see Table 1 in Methods) while maintaining the I_sptp_ and the cycle per pulse number as constants. When tuning tFUS parameters, we discovered that neuronal units grouped based on spike shape characteristics display different spike rate during the same tFUS stimulation. The excitatory neurons (RSUs) exhibit higher spike rates when stimulated with high UPRF, thus high duty cycle, whereas the inhibitory neurons (FSUs) exhibit high spike rate during stimulation at all UPRFs studied. The inhibitory phenomena found by Legon et al.(Legon et al., 2018; Legon et al., 2014) resulted from a UDC located in a transition zone between tFUS induction of inhibitory and excitatory effects (Plaksin et al., 2016), whereas the brain activation reported by Lee et al.(Lee et al., 2016a; Lee et al., 2016b) is probably due to the applied UPRF-related higher UDC. In other words, the UDC of 50% has already significantly increased the activity of excitatory neurons, and since the spiking activity of the inhibitory neurons does not increase proportionally, resulting in facilitation of behavioral outcomes.

### UPRF: Possible Mechanism of Cell-type Specific Effects

The UPRF, one type of modulation on the ultrasound temporal wave, may lead to a profound dynamic acoustic radiation force (ARF) (Renzhiglova et al., 2010). In a recent study, the ARF has been inferred as the most probable energy form that induces UPRF-dependent behavioral responses (Kubanek et al., 2018). In the present study, when different UPRFs are used to stimulate cortical neurons, we observed a significant difference in response between two neuron subpopulations. We also believe that the difference in response between different neuron types is observed due to the interactions between transcranial ARF and ion channels in the neuron membrane. Neurons exhibit different action potential waveforms due to the difference in distribution of membrane proteins both in channel types and relative quantity of each type of ion channel. These distinct types of membrane proteins may have different response dynamics to acoustic radiation force (Kubanek, 2018; Tyler, 2011). The basis of our hypothesis was demonstrated between the FSUs and RSUs in rat S1 cortex. An illustration of our hypothesis is shown in Fig. 1.

We further tested this hypothesis in the S1 cortex of transgenic mice. Optogenetics is used to identify excitatory neuron and inhibitory neuron populations, by coexpressing channelrhodopsin only in CamKIIa and PV expressing neurons. Although we cannot verify whether FSUs and RSUs correspond directly to PV neurons and CamKIIa neurons in this study, this model allows us to study specific protein expression to neuronal responses to tFUS stimulation (Fig. S7).

In this model, we also observed a distinct spiking response to tFUS stimulation UPRFs. Thus, in two different *in vivo* models and using two different methods to identify cell types, we have observed distinct responses to tFUS stimulation UPRFs. Based on our findings, future investigations can use genetic approaches to attribute the observed cell-type specific response to differences in protein expressions in different neuronal types.

### Sustained Synaptic Plasticity Effects of Ultrasound

In this long-term effect study, LTD was observed in 9 rats. Based on our hypothesis, we applied pulsed tFUS stimulation at 50-100 Hz stimulation in order to mimic the effects of LTP in the dentate gyrus. The observed results did not show LTP, but rather LTD, suggesting that the temporal encoding using tFUS does not share the same efficiency and/or mechanism as electrical tetanus stimulation. Additionally, the LTD induced by low-frequency electrical stimulation (LFS) is demonstrated (Fig. S8) to verify the brain location at the end of each animal experiment. However, the demonstrated long-term effect is a promising new feature of tFUS stimulation to be employed as a potential non-invasive therapeutic neuromodulation technique.

We also hypothesized that the UPRF will significantly affect the efficiency of LTD induction due to observed phenomenon in FSU and RSU cortical neurons. We did not see any clear trends supporting this hypothesis, possibly due to the differences in mechanisms studied in synaptic connectivity compared to individual neuron stimulation. Further investigations may still be needed to observe whether LTD strength and UPRFs are significantly correlated.

### Possible Mechanisms of Long Term Effects Observed from The Deep Brain

The frequency of electrical stimulation pulses results in increases below the action potential threshold in the postsynaptic potential, which in turn activates N-methyl-D-aspartate (NMDA) receptors and initiates a signaling cascade of transcriptions in the postsynaptic neuron (Luscher and Malenka, 2012). Several trains of high frequency tetanus electrical stimulations applied at 50 to 100 Hz for 1 second can induce LTP by causing a strong increase in presynaptic potential enough to activate the NMDA receptor, which in turn leads to a large increase in postsynaptic calcium concentration and initiation of signaling cascades. Similarly, low frequency tetanus electrical stimulation is applied at 1-3 Hz for several minutes in order to induce LTD. Here, the low frequency stimulation only leads to a modest increase in presynaptic potential, which is not enough to activate NMDA receptors but leads to prolonged subthreshold increase in postsynaptic calcium concentration due to repetitive stimulation, which triggers LTD (Sabatini et al., 2002).

In this study, the fact that LTP was not observed could be due to several factors. First, the mechanism of activation elicited by tFUS requires the delivery of repeated pulsed stimulation, which is more difficult to encode frequency dependent information in the presynaptic terminals. In contrast to electrical stimulation, where each electrical pulse applied at the presynaptic axon leads to a correlated increase in voltage, we may need to apply tFUS for multiple pulses in order to achieve the same effect. Also, due to the high duty cycles applied during LTD induction, there may be local temperature rises in focal spots of the ultrasound field. This is of particular concern because thermal changes have been known to modulate neural excitability and neurons will experience damage at significant deviations from physiological temperature ranges. No change in neural response as a function of temperature is expected, given that the estimated rise in temperature due to a 5-min exposure period of pulsed ultrasound (temporal sequence and spatial distribution are shown in Fig. 6B-E) with a UPRF of 3000 Hz, normal incidence and measured spatial-peak pressure amplitude of 99 kPa is only 0.43 °C when assuming no loss of heat (see Methods). Considering the profound skull heating when the focused ultrasound is propagating through such a bone structure, an estimated temperature rise at the skull-brain interface would be 0.97 °C. Due to the nature of single element focused ultrasound transducers in the axial focus, we cannot achieve focal activation only in the deep brain regions. Currently we cannot assert that in activating the deep brain, no other brain regions in the path of the ultrasound beam is activated, this would be achieved in future experiments using phased array focused ultrasound with refocusing techniques (Ballard et al., 2010).

### Ultrasound Safety

Except for ultrasound parameters employed in the subcortical region, all other tFUS stimulation parameters used on the S1 cortex of rats are maintained with I_spta_ below 50 mW/cm^2^, which lead to negligible temperature rises (< 0.001 °C) at the targeted brain area. In addition, the mechanical index (MI) used in these experiments is less than 0.1, given the low peak negative pressure (i.e. < 100 kPa). Such low MI makes cavitation in brain tissue unlikely. These levels are well within the levels advised by the Food and Drug Administration (FDA) standard for ultrasound diagnostic imaging safety (Duck, 2007; FDA, 2008). Hematoxylin and eosin stains gathered immediately after stimulation in both S1 and the hippocampus show no evidence of neuronal damage, local hemorrhage or inflammatory response at the stimulation site (see Fig. S5A-B and D).

### Study Limitation and Future Investigations

In this work, the rodent models were sedated by anesthetic agents, which may introduce an inevitable confounding factor of changing the neuronal spiking activities. In particular, the injection of ketamine/xylazine does not provide a constant anesthesia level. However, we have randomized the order of applied tFUS and Sham US conditions, which helps reduce the influence of anesthetic level to the statistical analyses. Based on literature reported first-order drug elimination kinetics, we established a numerical model to estimate the ketamine/xylazine blood concentration. When the anesthetic agents and ultrasound conditions were considered as factors, and the spike rates during the sonication were considered as responses, statistical tests indicate no significant effect in comparisons among ultrasound conditions by using the anesthetic agents (data not shown).

Caution should be taken when comparing between mice and rat models under tFUS stimulation. Our major constraint was due to the lack of widely available, well studied transgenic rat models for optogenetic stimulation. The thickness of the mice skull over the S1 cortex is 5 to 10 times thinner than that of rats. This leads to different distortions in tFUS field which may result in differences during stimulation. For mice subjects, a different collimator with a smaller tip size was used (see STAR Methods) to account for a smaller S1 region in order to avoid stimulating a widespread area in the mouse cortex. Discrepancies in activation area may also contribute to confounding results. Furthermore, rats were anesthetized with ketamine and xylazine cocktails while mice were anesthetized with isoflurane due to differences in experimental setup. Such a difference in anesthesia methods could also contribute to differences in results.

Regarding the effective harnessing of ultrasound energy, besides restricting the size of collimator outlet to be commensurate with the ultrasound wavelength, we are also cautious about the potential effect of using the 2-mm burr hole (Fig. 3B) via which the electrode array is able to reach the neurons. For this reason, we have introduced the 3D ultrasound field mappings (Fig. 2C-D; Fig. 6C-D) experimentally to explore potential alterations because of the low-acoustic-impedance conduit. And due to the requirement of 3D scanning of ultrasound field, we were only able to place the needle hydrophone (50 mm length) behind a freshly excised skull top piece in water, rather than inside a hollow rodent skull and conducted the volumetric pressure mapping. However, the latter one is believed to be more demanded so as to obtain additional knowledge of how significant of standing waves would be inside the rodent skull cavity by using the 500 kHz UFF, given that considerable interference patterns due to standing waves has been reported by administering 320 kHz tFUS to rats (Younan et al., 2013). The 3D computer simulations (Fig. 2E-F; Fig. 6E-F) were included to further depict and investigate the possible wave patterns inside the small skull structure while adapting the applied ultrasound parameters. To our knowledge, this is the first ultrasound field distribution simulation that takes into account the ultrasound propagation of the acoustic collimator.

Furthermore, as reported in (Sato et al., 2018; Tufail et al., 2011), the angles of the ultrasound incidence are designed to physically accommodate the ultrasound apparatus and the recording probe; however, we tried to preserve the ultrasound longitudinal wave rather than those in the shear mode. Although it is unavoidable in the practice, the angled tFUS may introduce nontrivial shear wave propagating along the skull, and thus lead to increased skull conduction. Nevertheless, the difference in the ultrasound wave mode might result in differed neuronal responses, which also necessitates further investigations.

### Potential Clinical Implications

The ability to selectively stimulate neural subpopulations non-invasively can provide a powerful scientific or clinical tool for disease models. For example, tFUS may be used to modulate atrophied brain regions in patients with Alzheimer disease to prevent disease progression or improve cognitive function. Studies have shown the Papez circuit in the anterior nucleus of the thalamus projects to multiple areas of the brain involving memory such as the dentate gyrus, anterior cingulate cortex and frontal and temporal regions. Deep brain stimulation of these regions has been explored to help improve memory (Fisher et al., 2010; Hamani et al., 2010; Hartikainen et al., 2014). We propose the cell-type selective effect of tFUS can be leveraged in conjunction with the existing phased array ultrasound technology to provide multi-focal tFUS stimulation (Hertzberg et al., 2010) to simultaneously target specific subset of neurons at different nodes of the brain memory circuit. This method will help achieve higher spatial specificity by ensuring no peripheral brain regions will be activated using ultrasound other than the regions of interest.

Furthermore, the ability to induce long-term synaptic depression may be used as a therapy for diseases involving hyperactivity or hypersynchrony of pathological neural tissues. In the case of Parkinson’s disease, pathological synchronization in the STN at the β frequency band, which can be disrupted by DBS stimulation (Eusebio et al., 2011), can presumably be altered through the application of noisy signals to perturb the local synchronized activity in the STN (Brown and Eusebio, 2008; Gatev et al., 2006; Hammond et al., 2007; Uhlhaas and Singer, 2006). Based on our results, we speculate tFUS can be used to perturb the synchronization in deep brain regions with potential for long lasting effects. Thus, tFUS can be a tool that patients use as an intermittent neuromodulatory therapy throughout the day.

## STAR METHODS

### Contact for Reagent and Resource Sharing

| REAGENT or RESOURCE | SOURCE | IDENTIFIER |
| --- | --- | --- |
| **Equipment** |  |  |
| **Single element focused ultrasound transducer** | Olympus Scientific Solutions Americas Inc. | V391-SU-F1.5IN-PTF |
| **Intracranial recording electrodes** | NeuroNexus | A1x32-Poly3-10mm-50-177 |
| **Recording system** | NeuroNexus | Smartbox |
| **Bipolar stimulation electrodes** | World Precision Instruments | TST33C40KT |
| **RF power amplifier** | Empower RF Systems | BBS0D3FHM |
| **Needle hydrophone** | Onda Corporation | HNR500 |
| **Chemicals** |  |  |
| **Furosemide** | Heritage Pharmaceuticals Inc. | NDC: 23155-473-44 |
| **Gentamicin** | VWR International | Cat#: 00149 |
| **Ketamine** | Zoetis Inc. | EA2489-564 |
| **Xylazine** | Akorn, Inc. | NDC: 59399-110-20 |
| **Bupivacaine** | Hospira/Pfizer, Inc. | NDC: 00409-1559-10 |

### Experimental Model and Subject Details

#### Rat Subjects

Wistar outbred male rats (Hsd:WI, Envigo, USA) were used as subjects, and all rat studies were approved by the Institutional Animal Care and Use Committee at University of Minnesota in accordance with US National Institutes of Health guidelines.

### Method Details

#### tFUS Setup and Parameter Selection

Single element focused transducers were used for tFUS stimulation. Transducer diameter 28.5 mm, ultrasound fundamental frequency (UFF) 0.5 MHz, −6dB bandwidth 300-690 kHz, a nominal focal distance of 38 mm. (V391-SU-F1.5IN-PTF, Olympus Scientific Solutions Americas, Inc., USA). Collimators were 3D printed with VeroClear material to match the focal length of the transducer and the animal model, the outlet of the angled collimator for the rat model has an elliptical area of 25.6 mm^2^ (major axis length: 6.8 mm, minor axis length: 5 mm), and the one for the ultrasound normal incidence has a circular area of 19.64 mm^2^. The size of collimators’ outlet was set to be no less than or at least commensurate with one ultrasound wavelength (i.e. 3 mm in soft tissue). One single-channel waveform generator (33220A, Keysight Technologies, Inc., USA) was working with another double-channel generator (33612A, Keysight Technologies, Inc., USA) to control the timing of each sonication, synchronize the ultrasound transmission with neural recording, and form the initial ultrasound waveform to be amplified, thus driving the transducer. A 50-watt wide-band radio-frequency (RF) power amplifier (BBS0D3FHM, Empower RF Systems, Inc., USA) was employed to amplify the low-voltage ultrasound waveform signal. The employed ultrasound intensity levels and duty cycles are described in Table. 1. As noted in the table, all ultrasound conditions used the same UFF of 0.5 MHz, ultrasound duration (UD, also known as sonication duration) of 67 ms, inter-sonication interval (ISoI) of 2.5 s, tone-burst duration (TBD) of 200 μs.

#### Extracellular Recordings

Extracellular recordings were made using 32-channel 10-mm single shank electrodes, where electrode sites are arranged in 3 columns, spaced 50 microns apart from each other (A1x32-Poly3-10mm-50-177, NeuroNexus, Ann Arbor, MI, USA). Electrodes were inserted into the rodent skull using a small animal stereotaxic frame with 10-micron precision manipulators (Model 963, David Kopf Instruments, Tujunga, CA, USA). Electrodes were inserted at a 40-degree angle in the sagittal plane, in order to directly record from brain areas under peak ultrasound stimulation.

Rodents were sedated under either ketamine and xylazine cocktail or isoflurane. All animals were skin prepared with hair shaving and hair removal gel. Rodent heart rate, respiration rates and inter-rectal temperatures were monitored throughout recording. Cranial windows, 1-2 mm in diameter, are opened in the skull using a high-speed micro drill (Model 1474, David Kopf Instruments, Tujunga, CA, USA) under stereotaxic surgery assisted with microscope system (V-series otology microscope, JEDMED, St. Louis, MO, USA). Brain suture lines were used to identify brain structure locations. Recordings were obtained using the NeuroNexus Smartbox recording system (20 kHz sample frequency, 16-bit ADC, NeuroNexus, Ann Arbor, MI, USA). All mentioned procedures on rodent models have been reviewed and approved by the Institutional Animal Care and Use Committee at University of Minnesota.

#### Stimulation at Primary Somatosensory Cortex in Rats

Rat subjects were adults in the weight range of 400 g to 600 g, sedated initially under ketamine and xylazine cocktail (75 mg/kg, 10 mg/kg) and extended with ketamine injections (75 mg/kg). This sedative approach allows us to achieve stable anesthesia with minimum body movement, e.g. breathing, heat beating, etc. tFUS collimator is coupled with the rat skull using ultrasound gel. Ultrasound collimators are transmitted through intact skull, directed at 40 degrees angle towards the left S1 region on the rat head.

#### Induction of Long Term Synaptic Changes in the Rat Hippocampus

Rats were initially sedated with ketamine and xylazine cocktail (75 mg/kg, 10 mg/kg), anesthesia extensions were done with Ketamine injections (75 mg/kg). Due to the increase pain level during surgery, local injections of Bupivacaine (5 mg/ml) were applied at the scalp. Bipolar tungsten electrodes (TST33C40KT, World Precision Instruments, USA) were inserted at 40 degrees into the medial perforant path (mPP) in order to stimulate the presynaptic terminal, a 16 or 32 channel recording electrode (NeuroNexus, Ann Arbor, MI, USA) was inserted at 50 degrees angle at the dentate gyrus (depth 3 mm) to record the resultant field excitatory postsynaptic potentials (fEPSP). fEPSP excitations were elicited with single square pulses with 200 μs to 1 ms pulse width and pulse amplitude of 0.7 to 1.7 V, delivered at frequencies between 0.4 to 0.1 Hz (PowerLab 26T, ADInstruments, Colorado Springs, CO, USA). Stimulation strengths were selected based on individual experiment setup and electrode location. The maximum energy delivered generating fEPSP without eliciting presynaptic action potentials was selected (O’Boyle et al., 2004).

A single-element focused transducer delivers tFUS stimulation at the dentate gyrus through the rat skull. Transducer interfaces with the skull via a collimator filled with ultrasound gel, with a tip diameter of 5 mm. 10-20 min of baseline fEPSP recorded before tFUS stimulation. Pulsed tFUS stimulation was delivered for 5 minutes at various UPRFs.

#### Auditory Brainstem Response (ABR) Test

ABR tests were performed on rats to determine the hearing threshold of rats before and after chemical deafening. Rodents were sedated under isoflurane (2%), 2 pairs of gold cup electrodes (Model 019-772500, Natus Medical Inc., Pleasanton, CA, USA) filled with EEG conductive paste (Ten20, Weaver and Company, Aurora, CO, USA) were applied to the rodent skull to record bilateral ABR (Akil et al., 2016). Each pair of electrodes consists of one reference electrode at the forehead immediately below the left or right eye, and on the ipsilateral side, a recording electrode at the base of the ear. A common ground electrode was placed at the base of the skull between both ears. Each electrode pairs were placed in differential mode between the recording electrode and the reference electrode to determine the brainstem response to a particular sound stimulus. Rodents were placed inside insulated sound chambers, auditory stimuli were aligned with the ear, tone bursts lasting 67 ms (same duration as our ultrasound stimulus) were pulsed at 5 Hz for 750 trial at each frequency and amplitude combination. Stimulus were delivered with a 5-cm round frame cone speaker (16 Ohm, frequency response: 0.5-10 kHz) (Model 289-131, Parts Express, USA), and its output frequency controlled by a waveform generator (33612A, Keysight Technologies, Inc., USA) spanning between 8 to 24 kHz, amplitude ranging between 56 to 92 dB.

To characterize whether neural responses observed are due to confounding effects of auditory stimulation, naïve animals received baseline ABR test prior to chemical deafening. Then, under isoflurane anesthesia, rats received tail vein injection of furosemide at 175 mg/kg (purchased from Boynton Pharmacy, University of Minnesota) followed by subcutaneous injection of gentamicin at 350 mg/kg (purchased from VWR International) (McGuinness and Shepherd, 2005). Post-injection ABR tests were performed 72 hours after initial injection. Once animals have been verified to have significant decrease in cochlear response, tFUS stimulation was applied at the somatosensory cortex, with concurrent intracranial recordings.

#### Histology

Explanted rodent brains were fixed in 5% formaldehyde for at least 48 hours and stored in 70% ethanol until samples were ready for histology. Samples were then dehydrated and embedded in paraffin for sectioned coronally at 5-10 micron per section. Tissue samples were then stained with Haemotoxylin and Eosin (H&E) by the Histology & Research Laboratory at the University of Minnesota. In some animals, magnetic nanoparticles (EMG308, Ferrotec Inc., USA) were injected via another type of NeuroNexus electrode array equipped with an injection duct (E16-20mm-100-177-D16, NeuroNexus, Ann Arbor, MI, USA) into the recording site after recording as a marker for the recording site. These samples followed the same fixation and sectioning procedures and the iron particles can be identified with Prussian Blue stain (see Supplementary Fig. S5C).

### Quantification Analysis

#### MUA Data Processing

For spike analysis, neural traces were band-passed between 244 Hz and 6 kHz, followed by Symlet wavelet denoising using Wavelet toolbox in MATLAB v9.0.0 (The MathWorks, Inc., Natick, MA, USA) to remove potential artifacts. All MUA spike sorting and single-unit preselection are performed using PCA based spike classification software Offline Sorter (Plexon, Dallas, TX, USA). Local field potentials (LFP) were band-passed from 1 Hz to 244 Hz, and denoised using Wiener filter and independent component analysis to generate Fig. 4E. Further analyses, including the ISpI computation, PSTH, raster and return plots (also known as Poincaré plot, a second-order analysis method for nonlinear features in time series), feature extraction for the initial phase (IP) and afterhyperpolarization (AHP), descriptive statistics for spike waveform and spike rates, LFP temporal and spectral analyses were performed using FieldTrip toolbox (Oostenveld et al., 2011) in the MATLAB. After obtaining phase durations of IP and AHP, K-means clustering function in the MATLAB was employed to conduct the cluster analysis of neuron types.

#### Ultrasound Pressure/Intensity Mapping

In order to characterize tFUS stimulation’s temporal and spatial dynamics near our targets, we developed a three-dimensional *ex-vivo* pressure mapping system that uses a water submerged needle hydrophone (HNR500, Onda Corporation, Sunnyvale, CA USA) driven by a 3-axial positioning stage (XSlide, Velmex, Inc., Bloomfield, NY, USA) to map out the spatial-temporal pressure profiles of ultrasound transmitted through an ex-vivo skull. The needle hydrophone was placed beneath an *ex-vivo* skull and recorded ultrasound pressure values at discrete locations (scanning resolution: 0.25 mm laterally, 0.5 mm axially) behind the skull. Skulls were freshly dissected from euthanized animals. This setup allows us to quantify the amount of energy delivered to the brain, which varies substantially due to the inhomogeneity and aperture of the skull. The system can be set up to mimic the exact conditions of the ultrasound set up with matched collimator locations and angles.

Based on the measured 3-D ultrasonic pressure map, the spatial-peak temporal-peak intensity (I_sptp_) is calculated using Eq. 1.

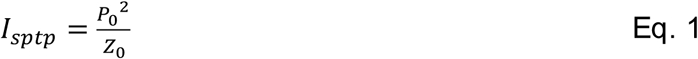

Where *P*_0_ is the maximal instantaneous pressure amplitude in both spatial and temporal domains, and *Z*_0_ is the characteristic acoustic impedance. This impedance can be computed using Eq. 2.

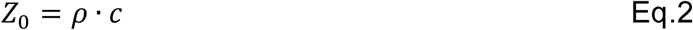

Where ρ is the medium density (1,028 kg/m^3^for brain tissue, 1,975 kg/m^3^for cortical bone (Culjat et al., 2010)), and c is the speed of sound in the medium (1,515 m/s for brain tissue, 3476 m/s for cortical bone (Culjat et al., 2010)).

We can also obtain the spatial-peak temporal-average intensity (I_spta_) from temporal profile of ultrasound pressure at its spatial maximum, which is denoted as *P*(*t*). Eq. 3 is used to calculate the I_spta_.

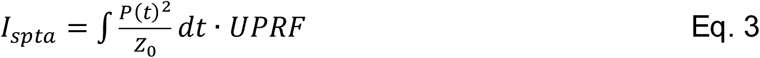

Moreover, the computer simulations to study the potential ultrasound wave patterns residing in the small skull cavity were conducted using a MATLAB-based toolbox, k-Wave (Treeby and Cox, 2010). The CT images of the Wistar rat were aquired from the scan using a Siemens Inveon microCT machine at the University Imaging Center of the University of Minnesota. The simulation study was following a similar protocol described in (Mueller et al., 2017) and using the acoustic parameters listed in Table 2. The skull was deemed to be immersed in water and its acoustic properties were mapped from the Hounsfield unit acquired from the CT images and thus the porosity readings. In the simulations, the Courant-Friedrichs-Lewy (CFL) number, the ratio of the wave travelling distance per time step to the space grid, was set at 0.02 to ensure stability of the simulations.

**Table 2.** Acoustic Parameters for Numerical Simulations.

| Materials | Speed of sound<br>(m/s) | Mass Density<br>(kg/m<sup>3</sup>) | Attenuation<br>Coefficient<br>(dB/MHz/cm) |
| --- | --- | --- | --- |
| Water | 1482 | 1000 | $3.02 \times 10^{-5}$ |
| Cranial Bone | 3100 | 2200 | Min: 1.87<br>Max: 18.15 |
| Collimator<br>(VeroClear™) | 2410 | 1160 | 9.54 |

#### Ultrasound Induced Temperature Rise

As a safety concern, once obtaining the ultrasound pressure map, we did a numerical estimation of maximum temperature rise using the following calculation methods (O’Brien, 2007). Firstly, we can obtain the I_spta_ from Eq. 3, and then the rate of heat generation per volume is calculated using Eq. 4.

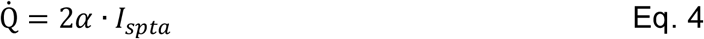

Where α is the ultrasound amplitude absorption coefficient in brain tissue (0.03 Np/cm at 0.5 MHz (Goss et al., 1978)) or in cortical bone (3.45 dB/cm at 0.5 MHz (Culjat et al., 2010)). Since we introduced the spatial-peak temporal-average intensity, the estimation of the Q value was maximized for the targeted site. Next, we borrowed the Eq. 5 from (Fry and Fry, 1953) to obtain the maximum temperature increase if we assume no heat removal process took place in the ultrasound energy deposition:

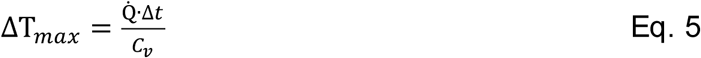

Where Δt is the tissue exposure time under tFUS, and C*_v_* is the heat capacity per unit volume (3.6 J/g/°C for the brain tissue, 1.606 J/g/°C for the skull (Ersen et al., 2014)). We estimated the temperature rise at a disk-shape focal area with a radius of one ultrasound wavelength (i.e. 3 mm). With these, the estimation would produce an upper limit for the temperature change.

### Statistical Methods

The animal or neuron numbers reflect our past experiences in developing neurotechnologies. Non-parametric statistical tests were conducted in R v3.2.1. The neuronal spiking data normality and variance homogeneity were initially inspected with Shapiro-Wilk test and Fligner-Killeen test, respectively.

#### Detailed Statistical Methods

##### *In-Vivo* Stimulation at Primary Somatosensory Cortex in Rats

###### Definition of center and dispersion

Neuron spike rate (Fig. 3F, G, I, and J): shown values are mean±95% confidence interval; Neuron spike rate (Fig. 5B, C, E, G, H): shown values are mean±s.e.m. Local field potential (LFP, Fig. 4E): shown values are mean±s.e.m.

###### Definition of n

Both number of animals and number of neurons as described in RESULTS.

###### Statistical test and definition of significance

For the spike rate analyses, significance (p < 0.05) was characterized by Kruskal-Wallis H test, and significance (p < 0.005) was characterized by *post hoc* one-tail two-sample Wilcoxon tests with Bonferroni correction for multiple comparisons.

For the LFP analyses, significance (p < 0.005) was characterized by permutation-based non-parametric statistics with false discovery rate (FDR) multiple-comparison correction.

###### Randomization strategy

The order of stimulation conditions (i.e., 30-4500 Hz UPRF tFUS stimulations, and corresponding sham ultrasound stimulation) was randomized.

###### Inclusion/exclusion of data

We included all possible single units based on the spike waveform, PSTH and inter-spike interval plots (time histograms and return plots).

Statistical details can be found in the captions of Fig. 3F, G, I and J, Fig. 4E, and Fig. 5B, C, E, G, and H in the Results sections of “Local Activations are Preserved in tFUS Setup When Auditory Pathway is Blocked”, and “Rat Somatosensory Cortex Exhibits Cell-type Specific Response to tFUS”.

##### Induction of Long Term Synaptic Changes in the Rat Hippocampus

###### Definition of center and dispersion

fEPSP slopes (Fig. 7A, C): shown values are mean±s.t.d.; % change from baseline slope (Fig. 7E): shown values are median with 25% and 75% quantiles.

###### Definition of n

Number of animals as described in RESULTS.

###### Statistical test and definition of significance

Significance (p < 0.001) was characterized by one-way ANOVA test with *post hoc* Tukey procedure.

###### Randomization strategy

The order of stimulation conditions (i.e., 3-10 kHz UPRF tFUS stimulations and sham stimulation) were randomized.

###### Inclusion/exclusion of data

We included fEPSP data that have distinct slope.

Statistical details can be found in the captions of Fig. 7A, C, and E in the Results section “tFUS Induces Long Term Changes in Synaptic Connectivity”.

##### tFUS Treatment Leads to Reduced Anesthesia Duration

###### Definition of center and dispersion

Anesthesia durations (Fig. S6D): shown values are mean±s.e.m.

###### Definition of n

Number of animals as described in the supplementary materials.

###### Statistical test and definition of significance

Significance (p < 0.05) was characterized by one-tail Wilcoxon test.

###### Randomization strategy

Each naïve rat subject was treated by only one condition among tFUS, SSKF or electrical stimulation only.

###### Inclusion/exclusion of data

We excluded failed anesthesia cases.

Statistical details can be found in the caption of Fig. S6D.

## SUPPLEMENTARY INFORMATION

Supplementary Information includes eight figures.

**Figure S1.** Computer Simulations for Mapping The Ultrasound Pressure Field inside The Skull Cavity, Related to Fig 2, Fig 6, Discussion and Methods.

**Figure S2.** Spatial Specificity of tFUS Induced Brain Activations, Related to Fig. 2.

**Figure S3.** The Spiking Statistics of The Fast-spiking Units, Related to Fig. 3.

**Figure S4.** Temporal Dynamics of Neuronal Action Potentials Responding to Administered Ultrasound and Sham Conditions, Related to Fig. 3.

**Figure S5.** tFUS Does Not Induce Neuron Damage, Local Hemorrhage or Inflammation, Related to Discussion and Methods.

**Figure S6.** tFUS Treatment Leads to Reduced Anesthesia Duration, Related to Discussion.

**Figure S7.** Validation of UPRF Preferences by Inhibitory and Excitatory Neurons, Related to Fig. 5 and Discussion.

**Figure S8.** Long Term Depression Induced by Low Frequency Electrical Stimulation (LFS), Related to Fig. 7.

## ACKNOWLEDGEMENTS

This work was supported in part by NIH grants MH114233, AT009263, EB021027, NS096761, and NSF grants CBET-1450956 and CBET-1264782. K.Y. was supported in part by an MnDRIVE Neuromodulation Fellowship and Doctoral Dissertation Fellowship from the University of Minnesota. X.N. was supported in part by Liang Ji Dian Graduate Fellowship at Carnegie Mellon University.

We thank Dr. Akira Sumiyoshi for providing Wistar rat MRI atlas, Dr. Abbas Sohrabpour for assisting EEG-based source imaging, Dr. Haiteng Jiang for inspiring discussion, Dr. Qi Shao for coordinating histology studies, Dr. Yi Zhang for training in animal surgery, John Basile for assisting equipment setup, Daniel Suma, Maryam Zhian, Mckinney Zhang and other members of the Biomedical Functional Imaging and Neuroengineering Lab for assistance.

## AUTHOR CONTRIBUTIONS

Conceptualization, K.Y., X.N., and B.H.; Methodology, K.Y., X.N., and E. K.-M.; Formal Analysis, K.Y.; Investigation, K.Y., X.N., and B.H.; Resources, B. H., and E.K.-M.; Writing – Original Draft, K.Y. and X.N.; Writing – Review & Editing, K.Y., X.N., E.K.-M. and B.H.; Supervision, B.H.

## DECLARATION OF INTERESTS

The authors declare no competing interests.

## DATA AND SOFTWARE AVAILABILITY

All data and software are available upon reasonable request to Bin He.

